# Approach-induced biases in human information sampling

**DOI:** 10.1101/047787

**Authors:** Laurence T. Hunt, Robb B. Rutledge, W. M. Nishantha Malalasekera, Steven W. Kennerley, Raymond J. Dolan

## Abstract

Information sampling is often biased towards seeking evidence that confirms one’s prior beliefs. Despite such biases being a pervasive feature of human behavior, their underlying causes remain unclear. Many accounts of these biases appeal to limitations of human hypothesis testing and cognition, de facto evoking notions of bounded rationality, but neglect more basic aspects of behavioral control. Here we demonstrate involvement of Pavlovian approach biases in determining which information humans will choose to sample. We collected a large novel dataset from 32,445 human subjects, making over 3 million decisions, who played a gambling task designed to measure the latent causes and extent of information-sampling biases. We identified three novel approach-related biases, formalized by comparing subject behavior to a dynamic programming model of optimal information gathering. These biases reflected the amount of information sampled (‘positive evidence approach’), the selection of which information to sample (‘sampling the favorite’), and the interaction between information sampling and subsequent choices (‘rejecting unsampled options’). The prevalence of all three biases was related to a Pavlovian approach-avoid parameter quantified within an entirely independent economic decision task. Our large dataset also revealed that individual differences in information seeking are a stable trait across multiple gameplays, and can be related to demographic measures including age and educational attainment. As well as revealing limitations in cognitive processing, our findings suggest information sampling biases reflect the expression of primitive, yet potentially ecologically adaptive, behavioral repertoires. One such behavior is sampling from options that will eventually be chosen, even when other sources of information are more pertinent for guiding future action.

## Introduction

Many spheres of human behavior depend upon gathering and understanding evidence appropriately to inform decision-making. Yet the best way to sample information is a nontrivial problem, necessitating deciding where to sample information (1, 2), when to cease information gathering (3, 4) and weighing up how such evidence should guide behavior (5, 6). Normative approaches can help address these questions (7), but their computational complexity renders them unlikely candidates for controlling behavior. Instead, these approaches can be better used as a basis for understanding limitations in cognitive processes and why biases emerge in human behavior (8, 9).

A particularly well-studied bias is that of confirming one’s prior beliefs (10). Inspired by classic rule discovery and falsification studies of Wason (11, 12), explanations of confirmation bias frequently appeal to limits in hypothesis testing as their latent cause. Several alternative accounts have been proposed. The ‘positive test account’ (13) posits that humans form beliefs about a particular hypothesis, and subsequently selectively seek and interpret evidence in support of this rule rather than against it. Yet it has been pointed out that this strategy may be normative in situations where possible competing hypotheses to explain the data are sparse (14). Other accounts suggest that humans are simply limited in the number of hypotheses they can consider at any given time (15).

It is widely acknowledged that humans are also subject to more primitive influences on behavioral control. Whilst these have been overlooked as a potential source of confirmation bias, they are known to impact upon information seeking in other domains. For instance, a primitive behavior present in several species is the observing response (16, 17). Here, animals select actions to yield information (reduce uncertainty) about the probability of receiving future reward, even when these actions have no bearing upon reward receipt. This can also be related to human preferences for revealing advance information about rewards when that information is immaterial to the task at hand (18). Critical here is the notion that in nature, advance information typically *is* valuable in guiding future action (unlike in the experimental tasks used to demonstrate these behaviors). Preferences for early temporal resolution of uncertainty (19) is thus conserved across humans and other species, and persists in influencing behavior even when rendered instrumentally irrelevant.

These considerations led us to consider how other primitive behaviors might bias information sampling. A notable characteristic of reward-guided behavior in many species is that of Pavlovian approach. Animals show greater efficacy in learning approach, as opposed to avoidance, actions that will lead to the delivery of reward (20, 21). Humans are also subject to similar approach biases (22). Pavlovian approach effects also spill over into the domain of attentional control, as stimuli previously ascribed a high value capture attention even when they are contextually irrelevant (23). As the locus of attention is intimately linked to information sampling during choice (24), this raises the possibility that Pavlovian approach may similarly influence information search.

To test this idea we examined gameplay data from a large-scale smartphone app (25) where we manipulated several factors of interest whilst probing subjects’ information sampling behavior. In brief, subjects played a card game in which they paid to sample information from different locations prior to deciding which option was most likely to yield reward. A framing manipulation meant that in half of all gameplays, approaching (choosing) the “biggest” option would be rewarded, but in the other half, approaching the “smallest” option would be rewarded. Crucially, the information structure of the task was identical across these matched conditions, such that any effects on information sampling could be ascribed to our manipulation as to the option subjects were instructed to approach.

We compared observed behavior to predictions derived from a normative dynamic programming model that computes the expected value associated with a perfect model of the task, treated as a Markov decision process (see methods and (26)). This enabled us to isolate three distinct biases in subjects’ information search that respectively influenced where information was sought, when information collection terminated, and how information was used to guide eventual choices. Relevant here is our recent quantification of human Pavlovian approach behavior parametrically in an approach-avoidance decision model on a separate economic decision task (27). We demonstrate that the prevalence of all three biases is related to this parameter.

## Results

### Information Seeking Task Design

Subjects played a binary choice game that involved paying escalating costs for information (by turning over playing cards), while gambling on which option was best based upon card values that were revealed (Fig 1A). There were six possible conditions that subjects might play (Fig 1B). Across three of these conditions, subjects’ objective was to identify the pair (row) of cards with the largest product (‘MULTIPLY BIGGEST’), largest sum (‘ADD BIGGEST’), or largest single card (‘FIND THE BIGGEST’). Across the remaining three conditions, the objective was inverted, such that they now sought the row with the smallest product, sum, or single card.

**Fig 1.**
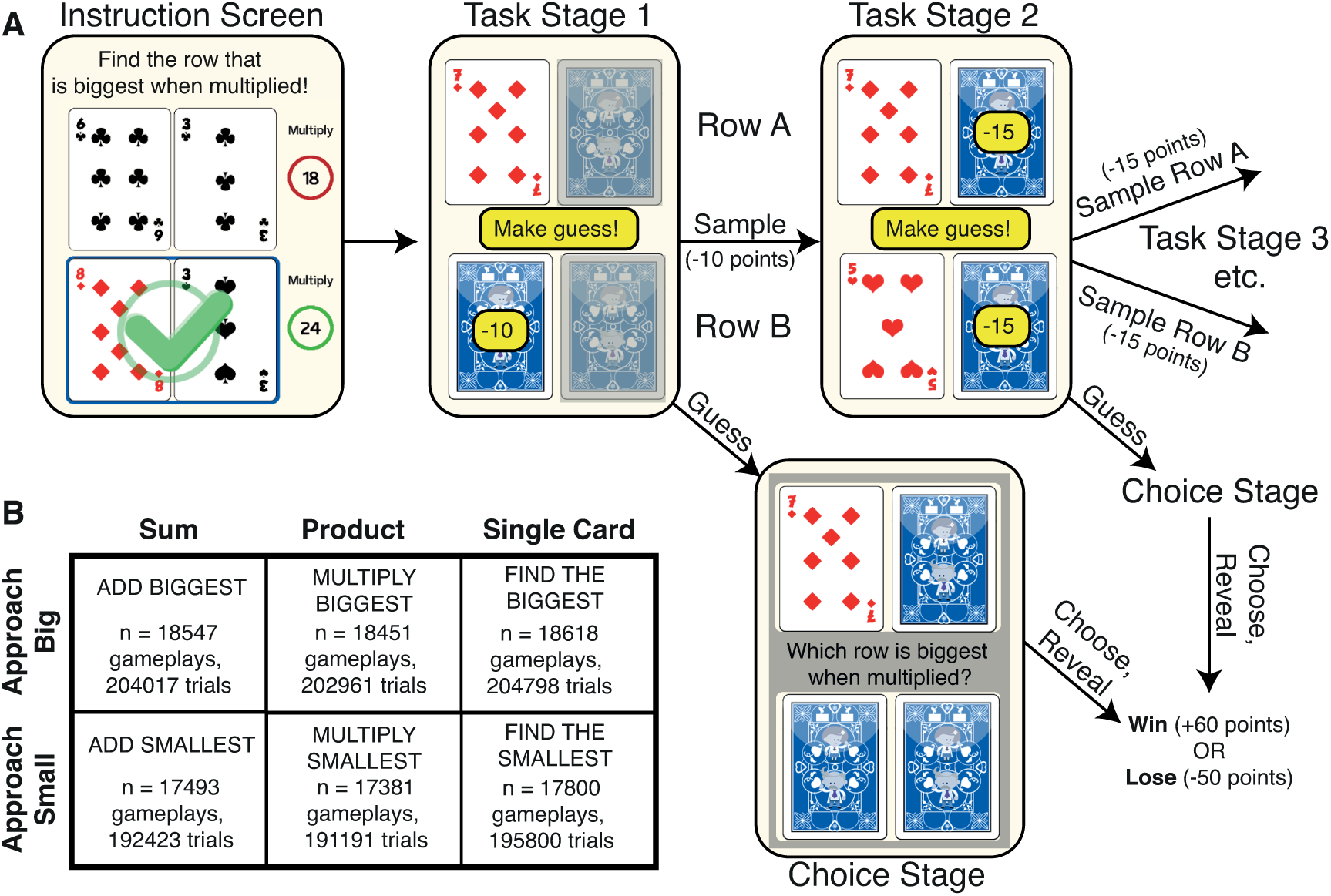
Information seeking task design. **(A)** Subjects aim to select the ‘winning row’ (in this example, the row with the largest product). After the first card is revealed (here, the 7 of diamonds), subjects enter task stage 1. Here they choose between the two yellow options, either sampling another card (costing 10 points) or making a guess about which is the winning row (no cost). Greyed-out cards cannot yet be sampled. If choosing to sample, then task stage 2 is entered, where either remaining card may be sampled (costing 15 points) or the subject may again guess. In task stage 3, sampling the last remaining card costs 20 points. At any task stage, making a guess means that subjects enter the choice stage. Here, after choosing, all cards are revealed and the subjected either wins 60 points if correct, or loses 50 points if incorrect. **(B)** The six task conditions, in a 3-by-2 design. Subjects either select the row with biggest (or smallest) sum, the biggest (or smallest) product, or the biggest (or smallest) single card.

At the beginning of each trial, all cards start face down. Subjects then touch the first card (randomly located) to turn it over, for no cost. This enters Task Stage 1 (Fig 1A). One of the three remaining cards is made available to be sampled at a cost of 10 points, but subjects can alternatively make a guess (gamble on which option will be rewarded) at no cost. If they choose to sample, the value of the second card is revealed and they enter Task Stage 2. Either of the two remaining cards can then be sampled at a cost of 15 points, or subjects can again choose to make a guess at no cost. If they choose to sample again, they enter Task Stage 3. The last remaining card can be sampled at a cost of 20 points, or they may again guess at no cost. At any Task Stage, making a guess means that subjects enter the Choice Stage. Here subjects choose which row they think will be rewarded, and all remaining cards are then turned face up. The subject wins 60 points if the gamble is correct, and loses 50 points if incorrect, minus the points paid for information sampling. Card values ranged, with a uniform distribution (sampled with replacement), from 1 to 10, with ‘picture cards’ removed from the deck.

On each gameplay, subjects were randomly assigned to play two short blocks (11 trials each) of two from the six possible conditions. The symmetry between the ‘approach big’ and ‘approach small’ conditions is crucial to our experimental design. Revealing a card of a particular value yields the same information content in both versions of the task (with the exception of the FIND THE BIGGEST and FIND THE SMALLEST conditions). This means that subjects’ information gathering behavior should, normatively, be matched across these conditions. The only behavior that should change is the final gamble made by the subject, which should reverse. By comparing across ADD BIG and ADD SMALL conditions, and across MULTIPLY BIG and MULTIPLY SMALL conditions, we could probe the influence of the approach direction (i.e., big/small) on information sampling behavior, and vice versa.

### ‘Positive evidence approach’bias

The first question we asked pertained to Task Stage 1 (Fig 1A). Here subjects decided whether to sample or guess based upon two variables: the information seen, i.e. the card value, and also the location where information was made available for sampling. We label the first row sampled as ‘row A’. In some trials, subjects were constrained to sample the next card from row A (‘AA trials’), whilst in other trials they were constrained to sample from row B (‘AB trials’).

As can be seen from the optimal dynamic programming model (Fig 2A), the card value and (to a lesser extent) the trial type influences the relative expected value of choosing to guess versus choosing to sample. The U-shaped function of the graph reflects an intuition that high‐ or low-valued cards are informative about the correct option to approach, making it more valuable to guess early. Mid-valued cards, by contrast, provide less information and make it more valuable to sample more information. The differential influence of AA versus AB trials is because the potential reduction in uncertainty depends upon the information that has already been revealed. Intuitively, on a MULTIPLY TRIAL where a 1 has been revealed, then sampling from row A again yields little information relative to row B, as it is already known that row A will have a low value (between 1 and 10). On a MULTIPLY TRIAL where a 10 has been revealed, then sampling from row A yields more information than row B as it reduces the range of possible row A values from between 10 and 100 to an exact value.

**Fig 2.**
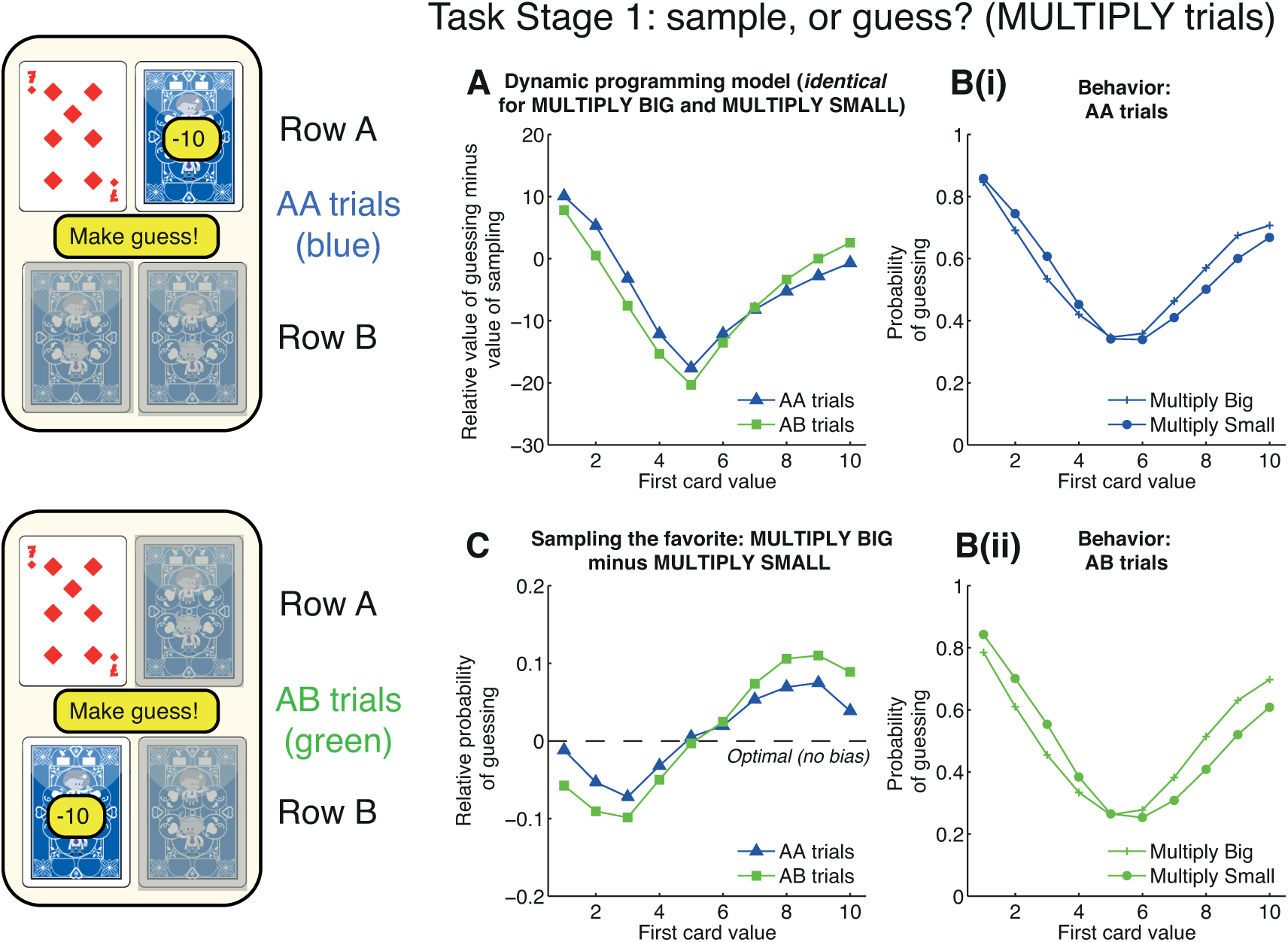
Positive evidence approach bias at task stage 1. At task stage 1, subjects decide whether to make a guess or pay 10 points to sample. The available card to sample may be on the same row (‘AA trials’) or the opposite row (‘AB trials’) as the first card. **(A)** Model predictions. The relative expected value (in points) of guessing vs. sampling from the dynamic programming model, in the MULTIPLY conditions. Mid-valued cards make it more valuable to sample, whereas extreme valued cards make it more valuable to guess. There is a weaker influence of the location of available information (compare ‘AA trials’ vs. ‘AB trials’). Crucially, optimal behaviour is identical for both MULTIPLY BIG and MULTIPLY SMALL conditions. **(B)** Subject behaviour. The probability of guessing in both conditions shows a broad similarity to the predictions of the dynamic programming model, but behaviour in MULTIPLY BIG and MULTIPLY SMALL shows systematic differences. (See Fig S1 for AA and AB trials plotted together, rather than MULTIPLY BIG and MULTIPLY SMALL.) **(C)** Positive evidence approach bias is revealed by subtracting the MUTLIPLY SMALL condition from the MULTIPLY BIG condition. Subjects are more likely to guess early if they have seen evidence that supports them approaching row A, rather than avoiding it. This effect is strengthened in AB trials, where subjects only have the opportunity to sample further information about row B. See also Figs S2/S3 for other conditions.

As expected, the dynamic programming model predicts identical behavior irrespective of the subject’s approach goal. As an example, consider revealing a 2 in the MULTIPLY BIG condition on an AA trial. This yields a probability of 0.764 that the ‘B’ row will be rewarded, and the expected value of guessing is therefore 0.764 *60 + (1-0.764)*(-50) = 34. The expected value of sampling again from the A row, calculated using dynamic programming, is 28.8, and so the relative expected value of guessing is 5.2 (Fig 2A). Now consider seeing a 2 in the MULTIPLY SMALL condition. This now yields the exact same probability of 0.764 that the ‘A’ row will be rewarded. Hence the expected value of guessing remains 34. The expected value of sampling further information remains 28.8, and so the relative expected value of guessing remains 5.2.

In contrast with these model predictions, subjects’ actual behavior showed a systematic difference between MULTIPLY BIG and MULTIPLY SMALL conditions (compare circles and plus signs in Fig 2B, see also Fig S1). Subjects became more likely to guess if they had seen evidence that supported them *approaching* row A rather than *avoiding* it. In MULTIPLY BIG, a high valued card (6 or above) carries evidence for choosing row A. Subjects become more likely to guess than when seeing the same card in MULTIPLY SMALL. However, a low valued card in MULTIPLY BIG (5 or below) carries evidence for avoiding row A. Subjects now become more likely to sample than when seeing the same card in MULTIPLY SMALL. This framing effect is seen most clearly when subtracting behavior in MULTIPLY SMALL from MULTIPLY BIG (Fig 2C).

The observed bias is one of approaching an option if positive evidence has been provided in support of that option, consequently we term this ‘positive evidence approach’. We observed positive evidence approach was more pronounced in AB trials than AA trials (Fig 2C). This is again consistent with our hypothesis, as subjects are less inclined to sample further information if available on a row they wish to avoid, than a row they wish to approach.

To quantify positive evidence approach across our population, we used the following summary statistic:

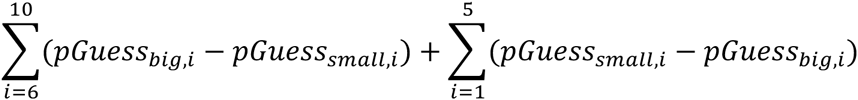

where *pGuess*_*big,i*_ and *pGuess*_*small*,*i*_ denote the average probability of guessing in MULTIPLY BIG and MULTIPLY SMALL respectively, having revealed card value *i*. As there should be no difference in the probability of guessing across the two conditions, the expected value of this statistic from the normative model is 0. By contrast, the value of this statistic across our population was 0.42 in AA trials, and 0.70 in AB trials. To estimate our confidence in this summary statistic, we recomputed it on 1,000 bootstrapped samples of 10,000 gameplays from our population. This yielded 95% confidence intervals of [0.30, 0.54] in AA trials, and [0.62, 0.79] in AB trials. (Throughout the paper, we focus on the reporting of effect sizes and 95% confidence intervals rather than p-values, as our large sample size renders p-values less informative (28)).

Similar results are seen by comparing the ADD BIG and ADD SMALL conditions (Fig S2; AA trials: mean 0.52, 95% CIs [0.40, 0.64]; AB trials: mean 0.79, 95% CIs [0.70, 0.89]). See also Fig S3 for FIND THE BIGGEST/FIND THE SMALLEST, which are not directly matched for information content.

It is also notable that overall, subjects’ behavioral choices to sample information were similar to predictions arising from the optimal model (Fig 2B), although not identical (Fig S1). This alone does not imply that subjects are implementing the optimal model. Instead, it may simply reflect the fact that relatively simple behavioral strategies will often recapitulate many features of more sophisticated strategies(29). For example, one straightforward strategy would be to compare the value of the presented card to the *average* value, estimate the current degree of uncertainty in making a choice, and then use these values with a softmax transformation (30) to calculate a probability for selecting row A, selecting row B, or sampling further information. We consider this question further in a latter section of the paper and show that this can approximate the average behavior of subjects in the task without recourse to an optimal model.

### ‘Rejecting unsampled options’ bias

We next asked how decisions to sample or reject information might influence subsequent choices. If subjects elected to guess at the first stage, they entered the Choice Stage, where they gambled on which option would be rewarded (Fig 1A). In ADD BIG, the relative expected value of choosing row A over row B increases with the value of the first card (Fig 3A, blue line), while in ADD SMALL it decreases with the value of the first card (Fig 3A, purple line). This was reflected in subjects’ choices in both sets of trials (Fig 3B; see Fig S4-S5 for other conditions). However, this decision arises on two different types of trial. The subject will either have just declined the opportunity of sampling information from the A row (on AA trials), or the B row (on AB trials). Our hypothesis was that information sampling depends upon the underlying approach value of an item. A corollary is that *declining* to sample an item reflects an underlying preference for *not* approaching it.

**Fig 3.**
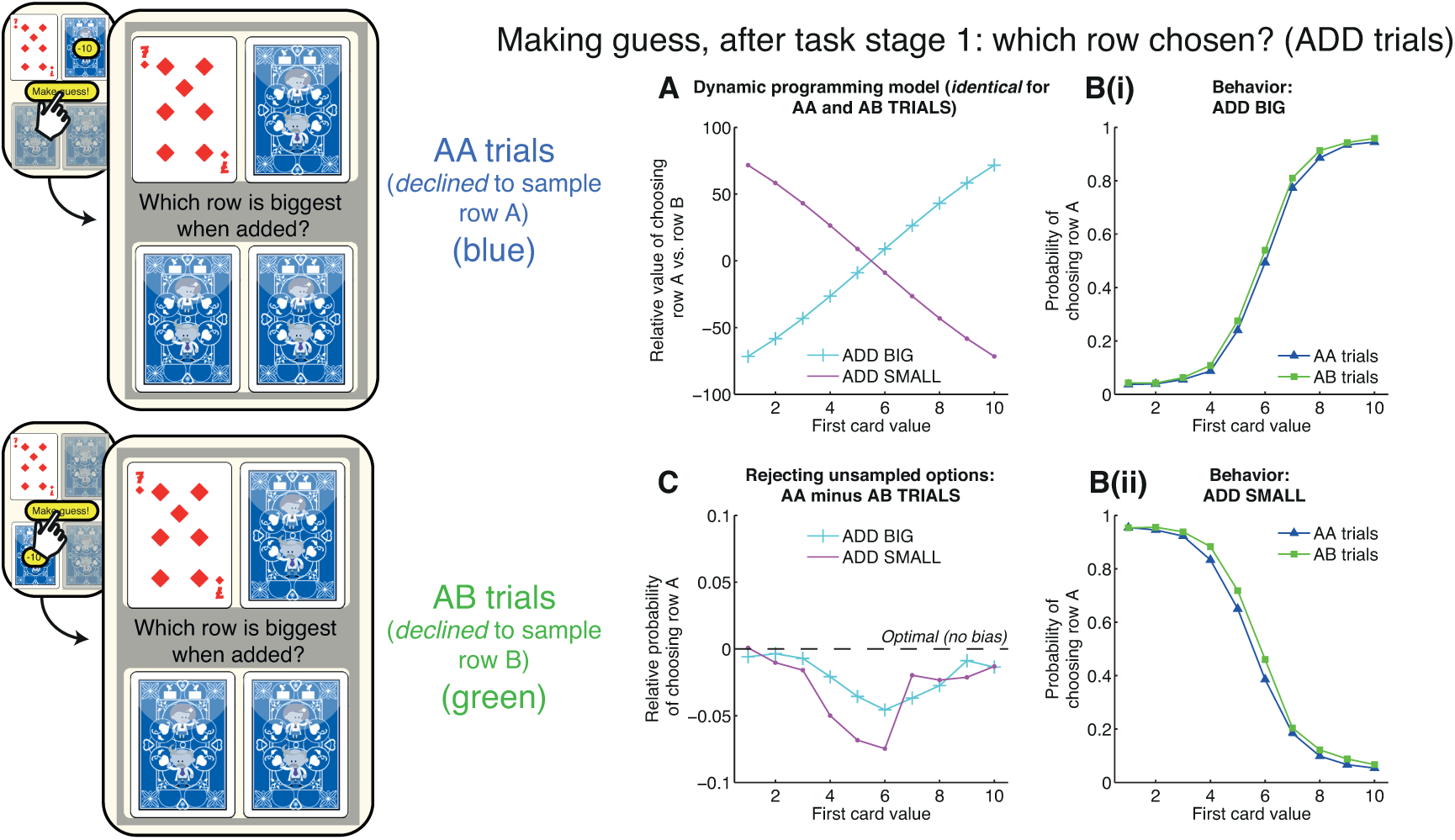
Rejecting unsampled option bias at choice stage. If subjects choose to guess at task stage 1, they then select between row A and row B. **(A)** Model predictions. The relative expected value (in points) of choosing row A vs. row B, in the ADD BIG (cyan) and ADD SMALL (purple) conditions. Crucially, the decision is identical for AA and AB trials; the only difference between these conditions is which row subjects have previously declined to sample. **(B)** Subject behaviour. The probability of choosing row A shows a softmax relationship to the relative values shown in Fig 3A. Note that the green line (AB trials) is higher than the blue line (AA trials) at nearly all card values, and particularly near the point of subjective equivalence. **(C)** Rejecting unsampled option bias is revealed by subtracting AB trials from AA trials, in both ADD BIG and ADD SMALL conditions. Subjects are more likely to choose row A if they have declined to sample row B. See also Figs S4/S5 for other conditions.

When we compared choice preferences on AA and AB trials for identical card values on the same condition we observed that, across all six conditions, subjects showed a systematic shift towards being less likely to choose the option that had just been left unsampled. Hence, subjects presented with the same card value on an AA trial were more likely to choose option B than on an equivalent AB trial (Fig 3B). This effect was most pronounced near the point of subjective equivalence in subjects’ choices, and is revealed most clearly by subtracting subjects’ choice behavior in AB from AA trials (Fig 3C).

We term this a ‘rejecting unsampled options’ bias. To quantify rejecting unsampled options across our population, we used the following summary statistic:

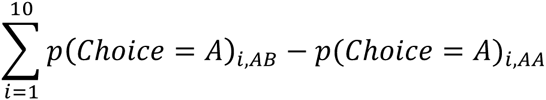

where *p(Choice = A)*_*i,AB*_ denotes the probability of choosing row A having observed card *i* on an AB trial, and *p(Choice = A)*_*i,AA*_ denotes the same probability on an equivalent AA trial. There is no difference in the choice that is presented to the subject between AB and AA trials, and the expected value of this statistic is therefore 0. The mean value of this statistic across the population was 0.21 in ADD BIG (95% CIs [0.10, 0.32]) and 0.30 in ADD SMALL (95% CIs [0.18, 0.41]).

We also found the rejecting unsampled options bias to be present across all the other conditions: MULTIPLY BIG (mean 0.24, 95% CIs [0.12, 0.35]; Fig S4), MULTIPLY SMALL (mean 0.27, 95% CIs [0.15, 0.39]; Fig S4), FIND THE BIGGEST (mean 0.22, 95% CIs [0.11, 0.35]; Fig S5) and FIND THE SMALLEST (mean 0.21, 95% CIs [0.09, 0.34]; Fig S5).

### ‘Sampling the favorite’ bias

Our design enabled us to also investigate where subjects chose to sample information. At Task Stage 2 on AB trials, we could determine subjects’ relative preference for sampling from row A versus row B (Fig 1A). Here, different conditions have different predictions for which row is more advantageous to sample. For example, in both the ADD BIG and ADD SMALL conditions, sampling from either row yields exactly the same amount of information about which row might be rewarded. The optimal dynamic programming model predicts no relative advantage for sampling from row A versus row B (Fig S6).

In both MULTIPLY BIG and MULTIPLY SMALL conditions, however, dynamic programming predicts that sampling from the row that currently has the higher-valued card will be more informative. The intuition behind this is that the range of possible outcomes on the row with the higher-valued card is greater, and so sampling further information on this row leads to a greater reduction in uncertainty than sampling the row with the lower-valued card. This is borne out in a heatmap of model predictions, showing the difference in relative value from sampling from row A versus row B (Fig 4A). Importantly, these predictions are identical for both MULTIPLY BIG and MULTIPLY SMALL conditions. Somewhat counterintuitively, it is therefore more advantageous to sample from the row with the largest card even in MULTIPLY SMALL. (Note that this is different from the relative expected value of guessing versus sampling, which is shown in Fig S8).

**Fig 4.**
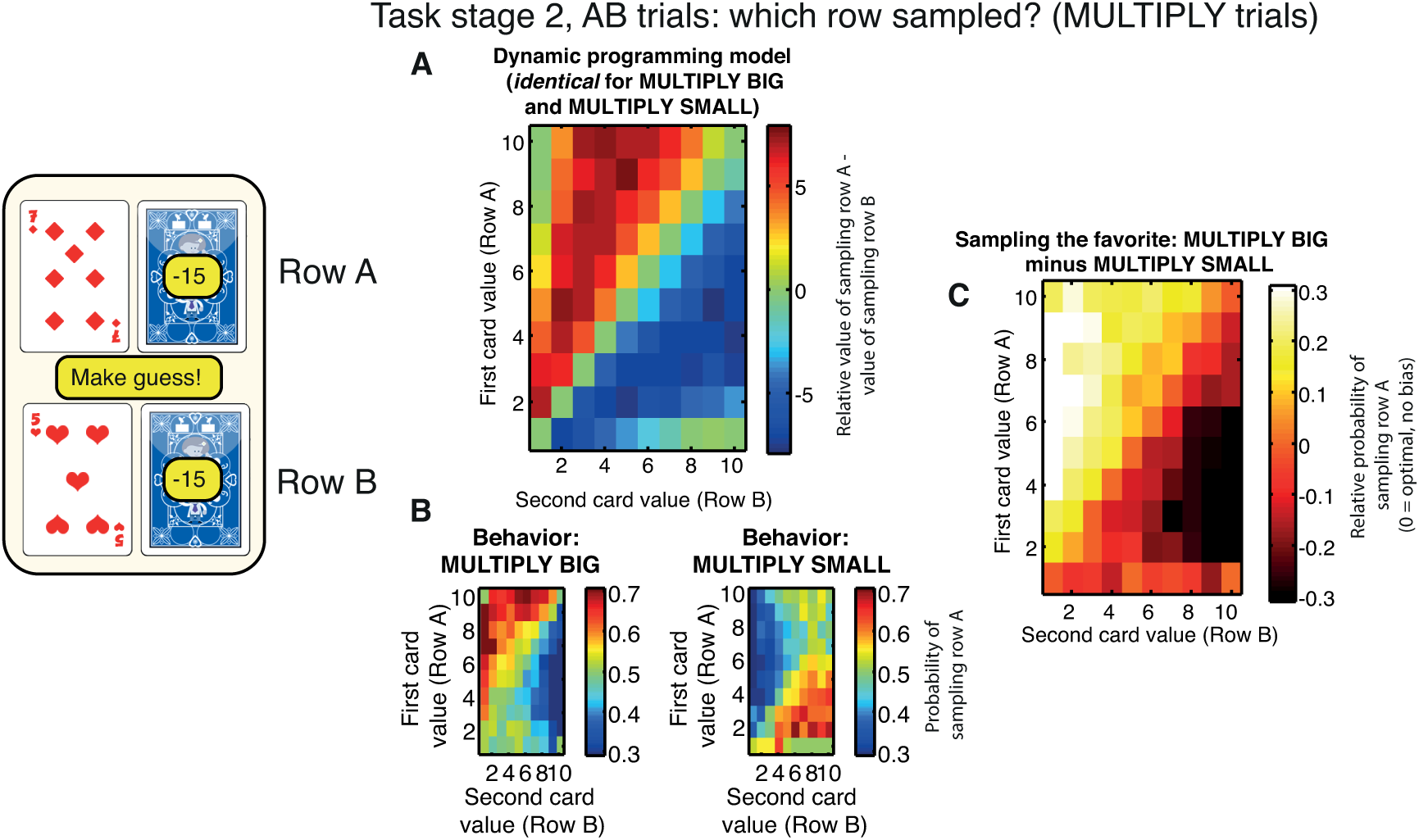
Sampling the favourite bias at task stage 2. If subjects choose to sample at task stage 1, they enter task stage 2. If this is on an AB trial, they then may sample again from row A, or from row B, or make a guess. **(A)** Model predictions. The relative expected value (in points) of sampling from row A vs. sampling from row B, in the MULTIPLY conditions. It is generally more valuable to sample from the row that currently has the higher valued card (e.g. the 7, in the example shown). Crucially, this prediction is the same in both MULTIPLY BIG and MULTIPLY SMALL conditions. **(B)** Subject behaviour. Subjects show a propensity to sample from the row containing the high-valued card in MULTIPLY BIG, but this trend is reversed in MULTIPLY SMALL. Subjects are therefore inclined to sample the option that is currently most likely to be approached. **(C)** Sampling the favourite bias is revealed by subtracting MULTIPLY SMALL trials from MULTIPLY BIG trials. The normative model predicts this heatmap to show no difference between conditions, yet there is a clear tendency to sample the currently favoured row. See also Figs S6/S7 for other conditions, and Fig S8 for relative value of guessing versus sampling across all six conditions.

In contrast to model predictions, we found that subjects preferred to sample from the option that currently had the higher value in the MULTIPLY BIG condition alone (Fig 4B, left). In the MULTIPLY SMALL condition, they preferred to sample from the option that currently had the lower value (Fig 4B, right). The influence of this bias in subjects’ information sampling is revealed more clearly by subtracting behavior in MULTIPLY SMALL from that of MULTIPLY BIG (Fig 4C). Whereas the optimal model shows no difference between these two conditions (i.e. the entire heatmap should equal 0), subjects reliably sampled information from the row that they currently sought to approach rather than avoid.

We term this bias ‘sampling the favorite’. To quantify sampling the favorite, we derived two statistics for trials in which subjects decided to sample a third piece of information. We calculated one ‘strong evidence’ statistic for trials in which the ‘favorite’ (the item that would eventually be approached) was clear. We defined this as trials where the difference in card values was 4 or greater in magnitude:

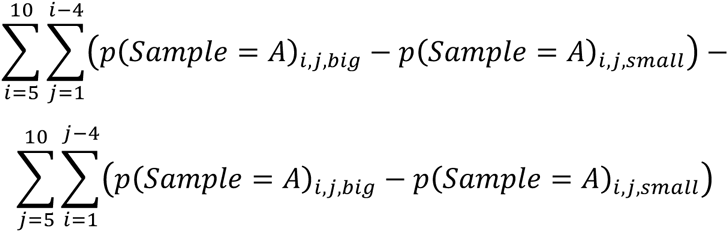

where *P(Sample = A)*_*i,j,big*_ refers to the relative probability of choosing to sample from row A over row B, when card *i* is presented on row A and card *j* presented on row B, on MULTIPLY BIG trials. The top row of the equation denotes trials where row A has a higher-valued card than row B, favoring approaching A in MULTIPLY BIG but approaching B in MULTIPLY SMALL. The converse is true for the bottom row.

We calculated a second ‘weak evidence’ statistic for trials in which the ‘favorite’ was less clear. We defined this as trials where the difference in card values was between 1 and 3 in magnitude:

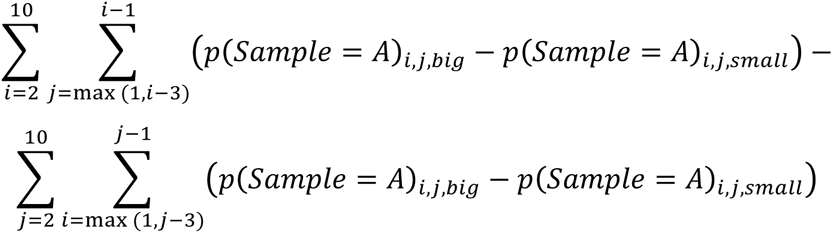

Crucially, because the optimal model predicts identical values for sampling from row A versus row B on MULTIPLY BIG and MULTIPLY SMALL, the expected value for both statistics is always 0. In contrast, the value of the ‘strong evidence’ statistic across our population was 12.35 (95% CIs [10.55, 14.01]), whilst the value of the ‘weak evidence’ statistic was 7.21 (95% CIs [5.85, 8.48]).

Note this bias was also observed in the ADD BIG/ADD SMALL condition (Fig S6), where the value of the ‘strong evidence’ statistic was 11.93 (95% CIs [10.39, 13.51]), whilst the value of the ‘weak evidence’ statistic was 6.86 (95% CIs [5.61, 8.14]). See Fig S7 for FIND THE BIGGEST/FIND THE SMALLEST, which are not directly matched for information content.

### A parametric model of subject behavior

We consider that subjects are unlikely to be implementing dynamic programming when they perform the task, yet their overall behavior shows a surprising resemblance to model predictions (e.g. Fig 2B, Fig 3B). We therefore constructed a simpler model that describes subjects’ performance without recourse to dynamic programming.

In this model, subjects first compute the value of choosing each option by comparing the presented value to the *average* value of all possible cards. In ADD BIG trials, at stage 1, for example, this would be:

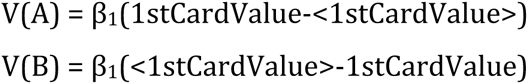

where <1stCardValue> is the expected value of the 1st card (5.5), and β_1_ is a free parameter. In ADD SMALL trials, we simply inverted the value of the each card, such that 10 became 1, 9 became 2, and so on.

We also considered an AB trial (e.g. Fig 4B), where the values of option A and B become:

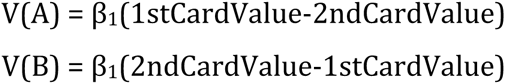

At both stages, we compute the degree of *uncertainty*, ω, in choosing either option:

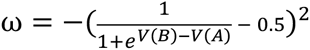

This is then used to derive the value of sampling information from option A or option B:

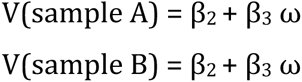

The probability of each action is finally generated using a softmax choice rule:

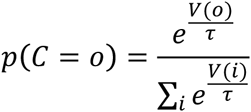

We fit parameters β_1_, β_2_, β_3_, and τ, using maximum likelihood estimation separately at stage 1 and stage 2. This model does not explicitly feature terms for the costs associated with sampling information; instead, these are implicitly factored into the constant term β_2_.

This model captures the main features of the behavioral data (Fig S9). At stage 1, it displays a U-shaped effect of card value on information sampling (as in Fig 2B) caused by the effects of choice uncertainty on the value of sampling information. It also displays a softmax choice curve (as in Fig 3B) between options A and B, matching subjects’ real choice probabilities between these two options. At stage 2, it displays choice probabilities between options A and B that again closely match subjects’ behavior.

However, because this model makes symmetric predictions for BIG and SMALL trials, it fails to capture the three biases described above (fig S9). We therefore adapted the model with three additional parameters to capture these biases (Fig 5). At stage 1, before entering values into the softmax choice equation, we captured the ‘rejecting unsampled options’ bias (Fig 5D-F) by adding an ‘approach bonus’ (β_4_) to the value of the item which could not be sampled. This makes subjects more likely to choose this option if they do not sample information.

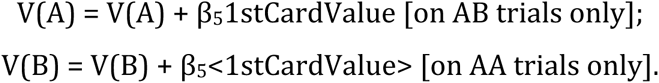

We also found that we could capture the ‘positive evidence approach’ bias (Fig 5A-C) by modulating the value of sampling option A:

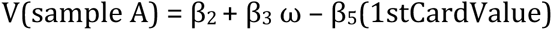

Notably, we found increasing 1^st^ card value had a negative influence on the value of sampling A, reflected by the negative sign in front of parameter β_5_. We infer that on AA trials (where option A is available to be sampled), subjects are more inclined to choose option A when it is of high value, than to sample again from it.

**Fig 5.**
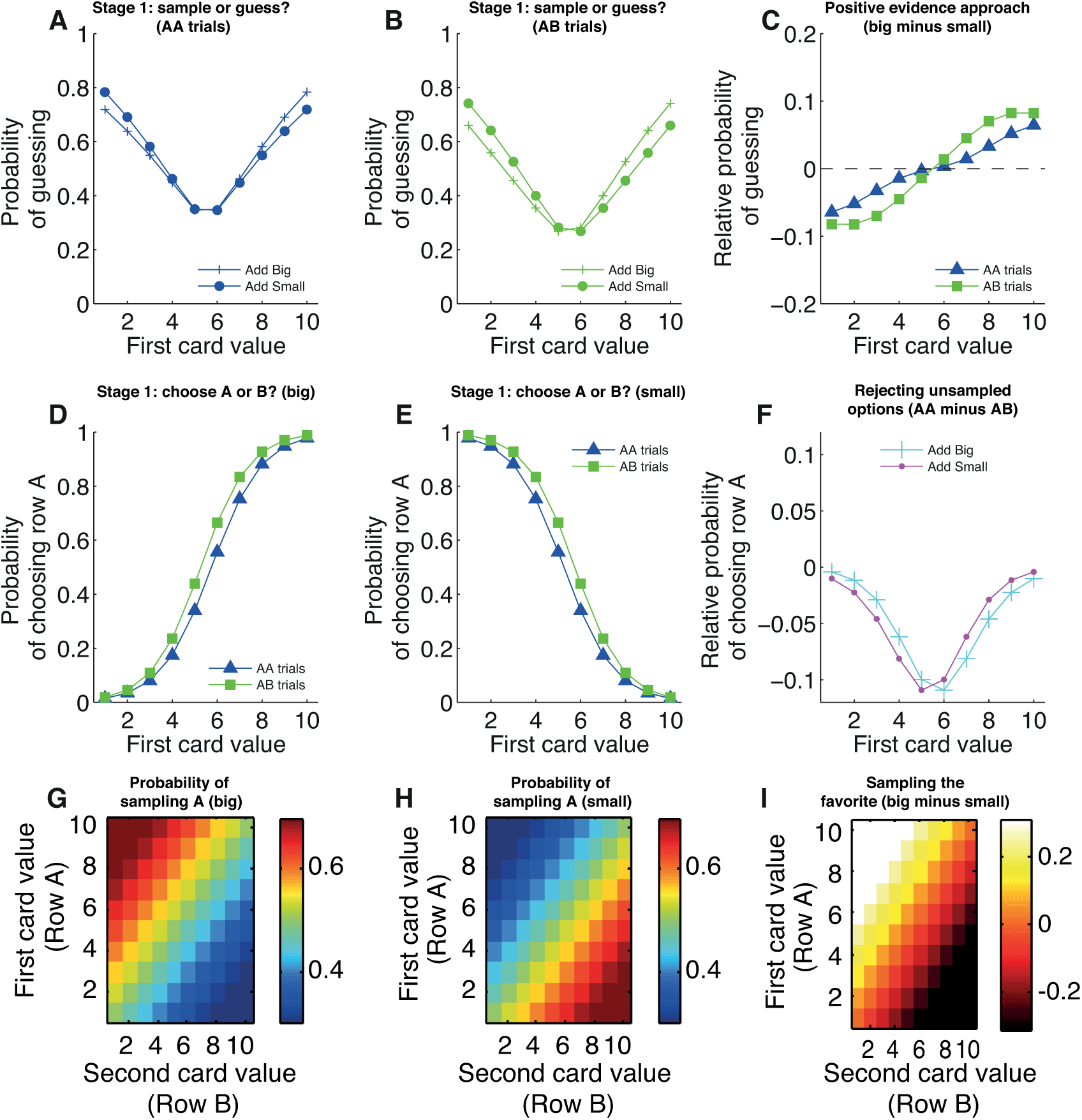
Behavioral predictions from the full parametric model of subject behavior, with best-fit parameters. **(A)** Predicted probability of guessing at stage 1 for AA trials (compare to Fig. 2Bi) and (B) AB trials (compare to Fig. 2Bii) in ‘add’ condition. **(C)** Predicted ‘positive evidence approach’ bias. Compare to fig. 2C. **(D)** Predicted probability of choosing row A, having chosen to guess, at stage 1, for ‘add big’ (compare to Fig. 3Bi) and (E) ‘add small’ (compare to Fig. 3Bii) conditions. **(F)** Predicted ‘rejecting unsampled options’ bias. Compare to Fig 3C. **(G)** Predicted probability of sampling row A in ‘big’ (compare to Fig 4Bi) and in (H) ‘small’ (compare to Fig 4Bii) conditions. **(I)** Predicted ‘sampling the favorite’ bias. Compare to Fig 4C. See also Fig S9.

Finally, at stage 2, we found that we could capture the ‘sampling the favorite’ bias (Fig 5G-I) by introducing a parameter that affected subjects’ propensity to sample from higher valued cards:

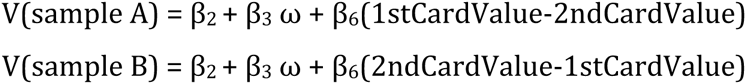

Parameter fits for stage 1 and stage 2 for both ADD and MULTIPLY trials are given in supplementary table S1.

The close fit between model predictions and subject behavior reveals that a far simpler framework (comparing a card value to the average expected value) can approximate an optimal dynamic programming model. Moreover, subjects’ approach-induced biases in information sampling can be readily parameterized within this framework. We anticipate that further, more refined models will be subsequently tested by downloading the raw behavioral data from DataDryad at http://dx.doi.org/10.5061/drvad.nb41c.

### Relationship of biases to Pavlovian approach-avoid parameter

An advantage of large-scale data collection via a smartphone app is that it allows data to be gathered on a range of cognitive tasks across a large cohort of subjects. Recently we reported learning and choice behavior on another gambling task contained within the same app platform (27, 31). In this simpler gambling task, subjects make binary choices between safe and risky options in three types of trials: gain trials (a certain gain versus a larger gain/zero gain gamble), mixed trials (certain zero gain versus a mixed gain/loss gamble), and loss trials (a certain loss versus a larger loss/zero loss gamble). Notably, subject behavior in this task was best characterized within a Pavlovian approach-avoidance decision model when compared to a range of models that also included a standard Prospect theory model (27). This decision model captures the influence on risk-taking behavior of both economic preferences and Pavlovian influences. It describes subjects’ value-independent propensity to approach or avoid gain gambles with a single parameter, *β*_*gain*_, and their value-independent propensity to approach or avoid loss gambles with a second parameter, *β*_*loss*_. Full details of modeling are provided in (27) and methods.

For each subject who played both games within the app (n=21866 users) we estimated *β*_*gain*_ and *β*_*loss*_ and computed the difference between these two parameters. We performed a median split on these values to derive two subpopulations of subjects, one exhibiting a larger bias for approach potential rewards over avoid potential losses, and one exhibiting a weaker bias. Next, we calculated the average behavior in our task of the subjects within these two subpopulations. We then fit the model described in the previous section to subjects’ aggregate behavior, and compared the fits of *β*_4_ (rejecting unsampled options), *β*_5_ (positive evidence approach) and *β*_6_ (sampling the favorite) statistics across the different subpopulations. To estimate our confidence in these statistics, we performed 100 bootstraps using 10,000 samples drawn from each subpopulation.

All three of our information sampling biases were differentially present in the high approach-avoid versus low approach-avoid groups. Positive evidence approach was greater in the high approach-avoid group in both add and multiply trials (Fig 6A). Rejecting the unsampled option was also greater in the high approach-avoid group in the add condition, although this difference was slightly reversed in the multiply condition (Fig 6B). Sampling the favorite showed a subtler pattern of expression was greater in the high approach-avoid group in both add and multiply trials (Fig 6C). All of the different observed biases are linked by the tendency to sample information from locations that will eventually be approached. The present results show that this is also reflected in the expression of these biases in groups exhibiting differential levels of Pavlovian approach influence on their behavior.

**Fig 6.**
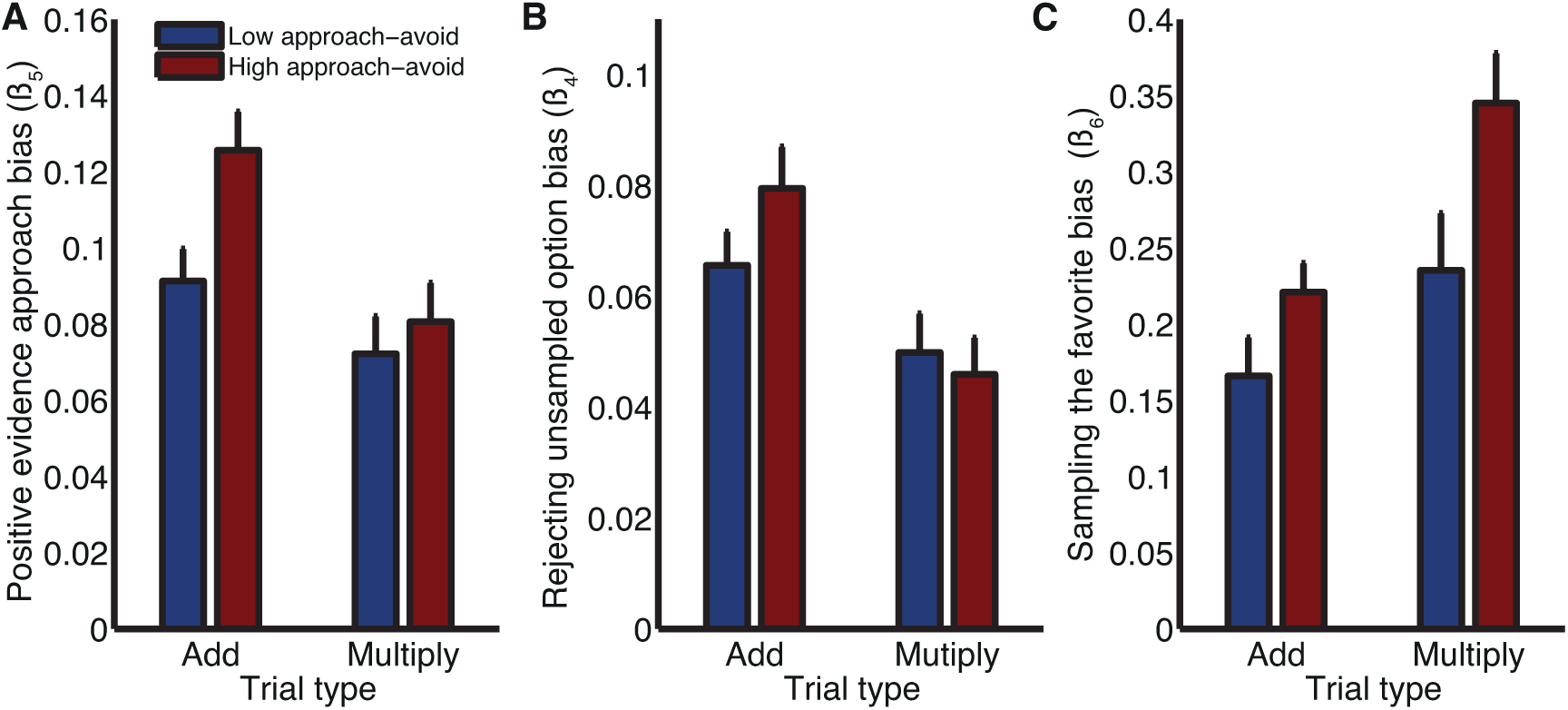
Expression of all three biases is differentially present in subjects with high vs. low Pavlovian approach, quantified in a separate gambling task. Blue bars denote subjects with below-median values for *β*_*gain*_ – *β*_*loss*_; red bars denote subjects with above-median values. **(A)** The positive evidence approach bias is quantified using the β_5_ parameter in the parametric model of subject behaviour; in both conditions, subjects with high approach-avoid parameter differences show more positive evidence approach than subjects with low parameter differences. **(B)** The rejecting unsampled option is quantified using the β_4_ parameter in the parametric model of subject behaviour; in the add conditions, there is considerably greater expression of rejecting unsampled option in subjects with high approach-avoid parameter differences; there is a weaker trend in the opposite direction in the multiply condition. **(C)** The sampling the favourite bias is by quantified using the β_6_ parameter in the parametric model of subject behaviour; in both conditions, subjects with high approach-avoid parameter differences show more sampling the favorite bias than subjects with low parameter differences. Bars/error bars reflect mean/s.d. across 1,000 bootstrapped samples of 10,000 gameplays.

### Variability in information sampling across age and education groups

An additional advantage of acquiring data via smartphone is that it enables examination of variation in information sampling across a much wider range of subjects than is typically examined in laboratory studies. In an initial exploration of this, we examined variation in a simple measure of information seeking, namely the average number of cards turned relative to the optimal model.

Subjects reliably sampled less information than predicted from the optimal model, but there was substantial variation across the population (Fig 7A). It is important to remember, however, that the model is only ‘optimal’ in the sense of maximizing expected points per gameplay. It does not, for example, include additional factors such as the subjective cost of sampling information. Indeed, we found that by adding a ‘subjective sampling cost’ of 5 points per turn to the optimal model shifted the distribution in Fig 7A so that it was now centred around zero (Fig S10). Nonetheless, variability in the extent to which individual subjects sampled information was highly reproducible across repeated gameplays (Fig 7B/S10), and we also found it to be stable irrespective of which set or ordering of conditions subjects played (Fig S11). This suggests that it provides a measure that might be related to performance on other cognitive tasks or demographic information about participants. An example of the latter is our finding that the number of cards gathered was positively related to both the highest level of attained education and age group of our participants (Fig 7C, top panels). Importantly, this measure was decoupled from general performance on the task, which was positively related to educational attainment but negatively related to age (Fig 7C, bottom panels). There was a very slight tendency for subjects with in the ‘high approach-avoid group’ to gather more evidence versus subjects in the ‘low approach-avoid group’, but this difference was negligible relative to the overall variance in information sampling across the population (mean of 0.0066 more cards sampled in high approach-avoid group, 95%CIs [0.0037, 0.0094]).

**Fig 7.**
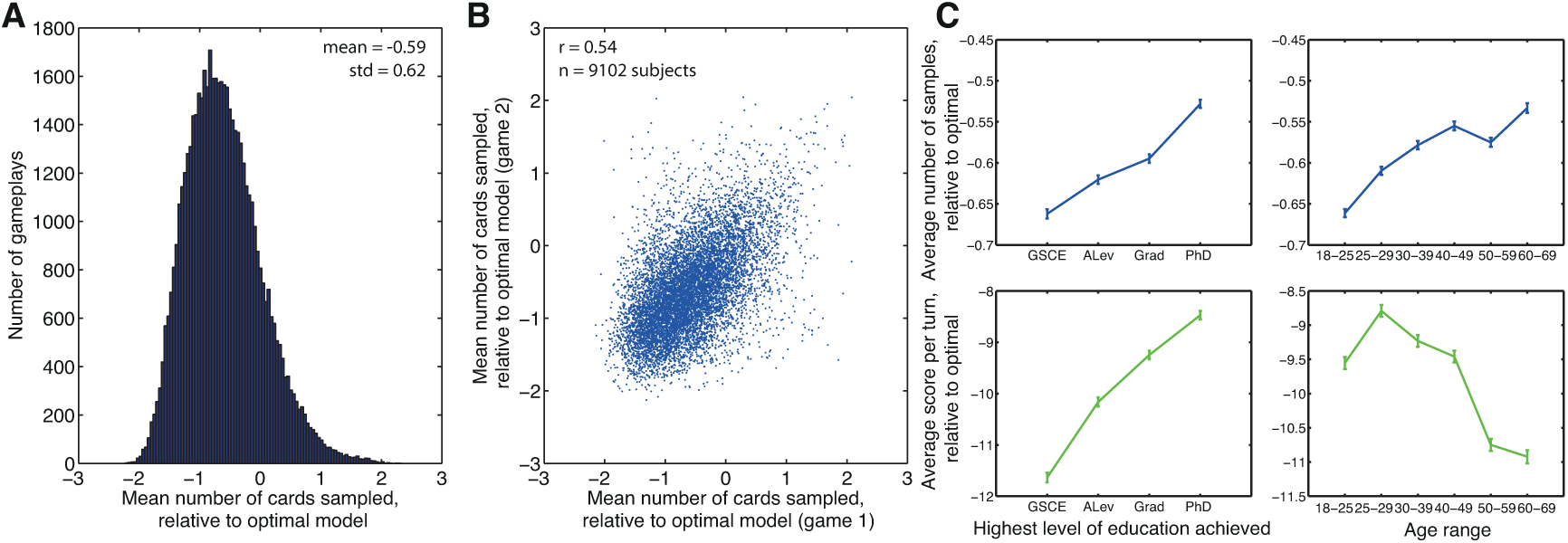
Individual differences in information seeking. **(A)** Histogram of the mean number of cards sampled in each trial, relative to how many would be sampled by the optimal model. Subjects show a propensity to guess early, but there is considerable individual variation. **(B)** Individual variation in information sampling reproduces across subsequent gameplays. Each dot is a subject; subjects who were inclined to seek little information in gameplay 1 also sought little information in gameplay 2. **(C)** Variation in information seeking (top) and subject performance (bottom) as a function of educational attainment (left panels) and age (right panels). General Certificate of Secondary Education (GCSE) is equivalent to 10^th^ Grade in United States; A Level (ALev) is equivalent to 12^th^ Grade. Datapoints denote mean +/- s.e.m.

## Discussion

Information seeking comprises interlinked decisions that includes how much to sample, where to sample from and finally which option to choose based upon sampled information. Whilst the complexity of our task allowed these different features to be indexed simultaneously within a single scenario, the task was sufficiently constrained that it can be treated as a Markov decision process. As such, an optimal model of the task can be derived using dynamic programming (26). Dynamic programming has rarely been considered as a normative basis for analysis of information search strategies in human information search (7). Although computationally expensive, a distinct advantage for our purposes is that it straightforwardly derives a common currency for the expected value of sampling in different locations against the value of choosing a particular option.

Subjects rapidly learnt the task, with their performance in terms of points gained becoming relatively stable within ~4 trials (data not shown); moreover, basic features of subject behavior (e.g. Figs 2B, 3B) matched with the overall pattern of predictions from the normative model. This confirms our previous observations concerning the validity of behavioral data acquired via smartphone (25). We make the large behavioral dataset freely available for download (via DataDryad, see http://dx.doi.org/10.5061/dryad.nb41c), providing an empirical testing ground for models of human information seeking.

Crucially, three features of subject behavior at different task stages showed demonstrable biases in information seeking. Two of these biases, positive evidence approach and sampling the favorite, were elicited as a consequence of our manipulation of which item subjects approached across different conditions. A third bias, rejecting unsampled options, was demonstrated as an effect of rejecting an option on the preference of a subject for choosing that option. All three biases were a consequence of the item that subjects currently sought to approach. Although manifesting as suboptimal biases in our experiment, we contend that these behaviors are present because they are likely to be, and have been, adaptive ecologically (32). In nature, foraging decisions (such as whether to stay or depart from a patch, or whether to engage with or reject an item of prey) are more common than those made between binary mutually exclusive options (33). In such contexts, we hypothesise that an adaptive strategy is to engage with the most valuable alternative first, and then accept or reject this alternative having acquired more information about its value. It would be intriguing to test whether approach-induced information sampling can produce optimal information sampling in more naturalistic foraging paradigms.

All three of our observed biases were differentially expressed in two groups who varied in the strength of expression of a Pavlovian approach-avoid parameter derived from a separate decision task. This provides a tentative suggestion of an underlying dopaminergic mechanism for control of Pavlovian approach on information seeking behaviors, given our recent demonstration that Pavlovian approach is boosted in subjects treated with L-DOPA (27). We also note that Polymorphisms in genes controlling dopamine function have recently been linked to individual differences in confirmation bias (34). Moreover, recordings from midbrain dopaminergic neurons reveal that they signal information in a manner consistent with the animal’s preference for advance information, in the same manner that they encode information about reward (17). Future studies could easily exploit possibilities of data collection via smartphone to test this and related hypotheses via combined collection of genetic and behavioral data across large populations. It might also be possible to design future versions of our task with a larger number of trials/conditions per subject, so as to elicit each of the three observed biases within-subject, rather than depending upon examining amalgamated data across a population.

It is possible that subjects had miscalibrated beliefs about task structure. For example, they may not have realized that there was a card value 1, which is normally replaced by an ‘ace’ in a regular deck of playing cards; or they may have believed that the average card value is 5, rather than 5.5. Such beliefs can straightforwardly be factored into the dynamic programming model, as can misunderstandings about the cost structure of the task, or additional opportunity costs for sampling further evidence. We found that such manipulations did indeed influence the relative preference of the model for guessing or sampling at different card values (not shown). Crucially, however, none of these belief-based manipulations predict any of the three biases observed. ‘Positive evidence approach’ and ‘sampling the favorite’ depend upon comparisons of SMALL and BIG conditions: *any* normative model predicts that subjects’ information sampling should be identical between these conditions, and that they should simply flip their final choice. Similarly, ‘rejecting unsampled options’ depends upon a comparison of final choice behavior in AA and AB trials, in situations where the subject has received identical information in both trial types.

It would also be possible to explore alternative versions of the current experiment that might examine the generality of our approach-avoidance account of information seeking biases. For instance, it would be intriguing to manipulate the affective valence of ‘points’ such that they became aversive, rather than rewarding. In such an experiment, we would predict that the approach-induced biases in information sampling would reverse. It would also be interesting to parametrically manipulate the costs involved in sampling different cards, as this would allow the experimenter to directly quantify the value of sampling information from different locations. It is also important to bear in mind that even when information sampling is biased, posterior beliefs can remain unbiased if belief updating is performed normatively (35, 36). It would be informative in future experiments to formally dissociate subjects’ apparent biases in information sampling from their biases, if any, in their belief updating.

Our findings are closely linked to other evidence from recent studies that relates the value of stimuli to deployment of attention (23, 37, 38). Both these studies, and our own, suggest that valuable items capture attention, and hence cause more information to be sampled from the associated location. In contrast with these previous studies, however, we show that the influence of value on information sampling occurs rapidly, can be reshaped depending upon current task goals, and can manifest as several distinct behavioral biases that affect multiple stages of information sampling. Combined, this evidence argues that choice models in which attention and information sampling are determined purely stochastically (6) require revision. Whereas these models convincingly demonstrate an important role for the locus of attention on valuation, the present data imply that the converse is also true. In simple terms, the value subjects ascribe to a location influences how likely they are to sample from it.

## Materials and Methods

### Smartphone-based data acquisition

Researchers at the Wellcome Trust Centre for Neuroimaging at University College London worked with White Bat Games to develop The Great Brain Experiment (25), available as a free download on iOS and Android systems (see http://thegreatbrainexperiment.com). Ethical approval for this study was obtained from University College London research ethics committee, application number 4354/001. On downloading the app, participants filled out a short demographic questionnaire and provided informed consent before proceeding to the games. Each time a participant started a game, a counter recording the number of plays was incremented. At completion of a game, if internet connectivity was available, a dataset was submitted to the server containing fields defining the game’s content and the responses given. The first time a participant completed any game the server assigned that device a unique ID number (UID). All further data submissions from that device, as well as the demographic information from the questionnaire, were linked to the UID. No personal identification of users was possible at any time.

The information-seeking game was available by clicking on ‘Am I a risk-taker?’, which launched the game. On each gameplay, subjects were randomly assigned to play short blocks (11 trials each, as outlined in Fig 1A and main text] of two different conditions randomly selected from six possibilities (Fig 1B). In two of these, subjects had to select the row that they believe contained the largest sum (‘ADD BIGGEST’] or largest product (‘MULTIPLY BIGGEST’]. In a further two conditions, subjects has to reverse their eventual choice and select the row containing the smallest sum (‘ADD SMALLEST’] or product (‘MULTIPLY SMALLEST’]. The remaining two conditions required participants to select the row with the largest or smallest individual card (‘FIND THE BIGGEST’ and ‘FIND THE SMALLEST’, respectively]. Full instructions for the task can be seen in Supplemental Text S1 and Movie S1. Raw data, along with MATLAB scripts for reproducing all figures shown in the paper, are made available for download on http://datadryad.org (doi:10.5061/dryad.nb41c].

The economic gambling task was available by clicking on ‘What makes me happy?’ Subjects started the game with 500 points and made 30 choices in each play. In each trial, subjects chose between a certain option and a gamble. Chosen gambles, represented as spinners, were resolved after a brief delay. Subjects were presented with the question, “How happy are you at this moment?” after every two to three trials.

### Dynamic Programming Model

The probabilistic structure of the task means that it is straightforward to derive a normative solution of task performance that maximises the expected average number of points to obtained from a given set of moves. This is achieved by applying dynamic programming to the task (30). At each step, dynamic programming calculates the expected value of every possible action (seeking more information in a particular location, or making a guess). To do so, it takes into account the full probability distribution of currently hidden cards, and the possible gain in information that can be obtained from sampling further.

Each combination of presented cards is defined as a state *s*. The best possible action *a* that a subject can take in a given state is defined as:

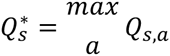

In a given state, the action value *Q* of making a particular guess in a particular state *s* can be calculated as:

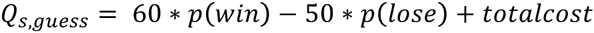

where *p(win)* is the current probability of winning by making that guess, *p(lose)* the probability of losing, and *totalcost* the incurred costs for sampling information thus far.

By contrast, sampling further information has a fixed probability (0.1) of transitioning into one of 10 possible subsequent states (10 different card values may be revealed). The value of sampling can then be calculated as the best action value in the subsequent state, multiplied by the probability of transitioning:

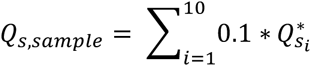

where si is the state that the subject would transition into if card value *i* is revealed. To calculate the value of sampling, one works backwards from the terminal state (all four cards revealed, where *Q*_*sguess*_ = 15 (=60-10-15-20)) to calculate 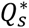 in all previous states. Full MATLAB model code is provided online at DataDryad, at http://dx.doi.org/10.5061/dryad.nb41c

### Pavlovian approach-avoidance model

Full details of the approach-avoidance decision model are given in reference (27). In brief, subjects’ expected utilities for choosing the safe option (*U*_*certain*_) and risky option (*U*_*gambie*_) were fitted using an established parametric decision model based on Prospect theory (39). The probability of choosing to gamble was then modelled by modifying the softmax function:

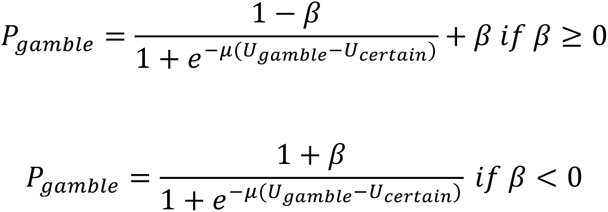

This causes a value-independent change in the probability of gambling, mapping choice probabilities to be bounded at (*β*,1) if *β* is greater than zero, and (0, *β*) if *β* is less than zero. *β* is fit separately for gain trials and loss trials, yielding two parameters, *β*_*gain*_ and *β*_*loss*_.

## Acknowledgments

We thank all members of The Great Brain Experiment team for support and interaction throughout the project, in particular Neil Millstone (White Bat Games) for designing and coding the app, and Peter Zeidman (Wellcome Trust Centre for Neuroimaging) for implementing server-side data collection and curation. L.T.H was supported by a Sir Henry Wellcome Postdoctoral Fellowship (098830/Z/12/Z). R.B.R. and R.J.D. were supported by the Max Planck Society. W.M.N.M. was supported by the Rosetrees’ Trust and the Astor Foundation. S.K. was supported by a Wellcome Trust New Investigator Award (096689/Z/11/Z), and R.J.D. by a Senior Investigator Award (098362/Z/12/Z). The Wellcome Trust Centre for Neuroimaging is supported by core funding from Wellcome Trust Grant 091593/Z/10/Z. Initial app development was funded by Wellcome Trust Engaging Science: Brain Awareness Week Award 101252/Z/13/Z. We also thank Jonathan Nelson and one further anonymous reviewer for their detailed and thoughtful comments on our manuscript during the review process.

## Supporting Information Figure legends

**Fig S1. Probability of guess in MULTIPLY condition for MULTIPLY BIG (left) and MULTIPLY SMALL (right) conditions**. Data are the same as in main figure 2B, but are replotted with AA and AB trials on top of each other to facilitate comparison with dynamic programming model predictions.

**Fig S2. Figure layout as per main Fig 2, but for ADD conditions**. Note that there is now no difference between normative model predictions for AA vs. AB trials.

**Fig S3. Figure layout as per main Fig 2, but for single card trials**. Note that in single card trials, the same card value carries different amounts of information between the two conditions (hence part A is split into two plots). As such, there is no direct equivalent for the positive evidence approach bias.

**Fig S4. Figure layout as per main Fig 3, but for multiply trials**.

**Fig S5. Figure layout as per main Fig 3, but for single card trials**.

**Fig S6. Figure layout as per main Fig 4, but for add trials**. Note that there is no relative advantage for sampling row A over row B in the normative model (part A).

**Fig S7. Figure layout as per main Fig 4, but for single card trials**. Note that in single card trials, the same card value carries different amounts of information between the two conditions (hence part A is split into two plots). As such, there is no direct equivalent for the sampling the favourite bias.

**Fig S8. The relative expected value of guessing (and choosing the best option) minus the expected value of sampling further information, at Task Stage 2 on AB trials, derived using dynamic programming**. (A) ADD conditions, where predictions are identical for ADD BIG and ADD SMALL. (B) MULTIPLY conditions, where predictions are identical for MULTIPLY BIG and MULTIPLY SMALL. (C) SINGLE CARD conditions. Top row = FIND THE BIGGEST condition, bottom row = FIND THE SMALLEST condition.

**Fig. S9. Behavioral predictions from the reduced (4-parameter) model of subject behavior, with best-fit parameters**. Data is plotted as in main Fig. 5. **(A)** Predicted probability of guessing at stage 1 for AA trials (compare to Fig. 2Bi) and **(B)** AB trials (compare to Fig. 2Bii) in ‘add’ condition. **(C)** Predicted ‘positive evidence approach’ bias. Compare to fig. 2C. **(D)** Predicted probability of choosing row A, having chosen to guess, at stage 1, for ‘add big’ (compare to Fig. 3Bi) and **(E)** ‘add small’ (compare to Fig. 3Bii) conditions. **(F)** Predicted ‘rejecting unsampled options’ bias. Compare to Fig 3C. (G) Predicted probability of sampling row A in ‘big’ (compare to Fig 4Bi) and in **(H)** ‘small’ (compare to Fig 4Bii) conditions. **(I)** Predicted ‘sampling the favorite’ bias. Compare to Fig 4C.

**Fig. S10. Data plotted as in main figure 7A/B, but with an additional ‘subjective sampling cost’ of 5 points/turn added to the normative model**. The mean of the distribution of the number of cards sampled relative to the model (left panel) now lies close to 0.

**Fig S11. Information seeking is a stable trait irrespective of condition ordering**. Along the bottom of the matrix is the condition experienced in the first 11 trials of gameplay 1 (1 = FIND BIGGEST, 2 = FIND SMALLEST, 3 = ADD BIG, 4 = ADD SMALL, 5 = MULTIPLY BIG, 6 = MULTIPLY SMALL), whilst along the left of the matrix is the condition experienced in the first 11 trials of gameplay 2. The color of the heatmap reflects the correlation coefficient between information sampling (relative to the optimal model) across the two gameplays (as in main Fig 7B).

## Supporting Information Table

**Table S1. Maximum likelihood estimates of parametric model of subject behavior.**

## Supporting Information Text

**Text S1. Instructions provided to subjects when performing the task.**

## Supporting Information Movie

**Movie S1. Example movie of subject performing several trials of the task, starting from the home screen**. (Readers can also download the app, at http://www.thegreatbrainexperiment.com).

